# Single cell sequencing of zebrafish kidney marrows reveals AHR2-dependent endogenous regulation of hematopoiesis

**DOI:** 10.1101/2024.04.23.590755

**Authors:** Subham Dasgupta, Britton Goodale, Robyn Tanguay

## Abstract

The aryl hydrocarbon receptor (AHR) is a ligand-dependent transcription factor that mediates a wide range of biological and toxicological responses. While largely studied in ligand-activated toxicant responses, AHR also plays important roles in endogenous physiological processes. We leveraged single cell sequencing and an AHR2 knockout zebrafish line to investigate the role of AHR2 in regulating hematopoiesis (production and differentiation of red and white blood cells from hematopoietic stem cells). Our objectives were to determine if absence of AHR2-1) alters proportions of immune cell populations and/or 2) impacts gene expression within individual immune cell types. We dissected kidney marrow (organ of hematopoiesis in zebrafish) from adult wildtype and AHR2 knockout zebrafish (N=3/genotype), isolated single cells and sequenced ∼ 5000 cells/sample (10X Genomics). We identified 14 cell clusters representing the expected major blood (erythrocytes, thrombocytes), immune (B cells, macrophages, lymphoid cells, granulocytes, etc), progenitors and kidney cell populations. We focused our analyses only on the progenitor and mature immune cell types. While there were no genotype-specific differences in proportion of individual cell types, gene expression differences were observed within several cell types. For known genes, such as *rrm2*, changes were up to 2000-fold, signifying their importance in AHR2-hematopoesis interaction. Several of the known genes are also identified as markers of carcinoma cells for an array of cancer types. However, many of the dysregulated genes are poorly annotated, limiting our ability to examine biological processes and pathways dysregulated on AHR2 mutation. Nevertheless, our study indicates that AHR2 plays an important endogenous role in hematopoiesis. Future work will focus on better characterizing anatomy of dysregulated genes and their functions in hematopoiesis.

## Introduction

The aryl hydrocarbon receptor (AHR) acts as a transcriptional regulator, binding to aromatic hydrocarbon-responsive elements (AHREs) and exerting physiological functions through ligand-binding, dimerization and interactions with other proteins. Toxic effects of environmental contaminants such as polycyclic aromatic hydrocarbons, 2,3,7,8-Tetrachlorodibenzo-p-dioxin (TCDD), polychlorinated biphenyls (PCBs) are widely known to be mediated by AHR ^1^. Indeed, investigation into contaminant-induced AHR-mediated immunotoxicity, reproductive toxicity, developmental toxicity and other toxicological mechanisms remain active areas of research^1^. However, the endogenous role of AHR in normal physiology remains less explored.

The current study uses an AHR2-null zebrafish line to study endogenous role of AHR2 in hematopoiesis, focusing on the immune system. In zebrafish, AHR2 is the primary functional orthologue of human AHR^1^. Since AHR-associated studies typically leverage AHR ligands to study mechanisms through AHR activation, the area of endogenous or basal AHR functions remains an area of active investigation. The process of hematopoiesis constitutes development of the blood and immune cells from hematopoietic stem cells (HSCs). The immune cells, in turn, constitute of a battery of cell types, each with its own role in establishing defense mechanisms^2^. Therefore, understanding its regulators is crucial from a basic physiology perspective. Prior studies have suggested that AHR has a basal role in regulation of lymphocytes and T cell function^3^; however, the evidence is not clear. Additionally, mechanistic assessments are largely conducted in *in vitro* models, failing to provide a holistic view of AHR regulation of multitudes of immune cell types. Within this study, we coupled single cell sequencing with AHR2-null zebrafish line to examine effects of AHR2 mutation on progenitor and mature immune cell populations and gene expression. This enabled us to profile the distribution of immune cell types and gene expression within each cell type as a function of AHR2 regulation.

## Methods

### Animal husbandry

Male tropical 5D wildtype (*ahr2*^*+*^) and AHR2 null (*ahr2*^osu1^) were obtained from Sinnhuber Aquatic Research Laboratory, Oregon State University and transported to Dartmouth College, Geisel School of Medicine. These fish were previously generated using CRISPR-Cas9 and well-characterized in our prior work^1^. Adult animals were raised in a recirculating water system (28 ± 1°C) with a 14-h:10-h light-dark schedule. Adult wild-type fish were kept at a density of 6–8 fish per liter. Adult were euthanized with Tricane (3-amino benzoic acid ethylester) at concentrations exceeding 300–400 mg/L prior to dissection of kidney.

### Kidney dissection and single cell extraction

Three male fish per genotype were used for our studies for robust biological replication, resulting in a total of 6 samples. Since we were limited by the number of fish we could transport, male fish were used to avoid confounding effects of presence/absence of egg clutches between female fish on the day of dissection. Head regions of the kidney, which contains the kidney marrow and constitutes the primary site of hematopoiesis in adult zebrafish were dissected from each fish. Care was taken to avoid inclusion of blood vessels as lysis of red blood cells can lead to contamination of ambient RNA with hemoglobin (Hb) genes in droplet-based single cell sequencing methods. Dissected kidneys were transferred into 2 ml Eppendorf tubes containing 500 µL of modified 1X phosphate buffer saline (PBS) supplemented with 0.5% bovine serum albumin and 1% Penicillin-Streptomycin. Single cell suspensions were obtained by dissociating tissues with 200 µL pipette tips and a 20-gauge needle, followed by washing with PBS and filtering through a 40 µm cell strainer (Corning). Cells were counted and viability assessed using a Luna II cell counter (Logos Biosystems) with trypan blue, and dead cells were removed using a Dead Cell Removal kit (StemCell).

### Single cell sequencing

Single cell suspensions of the 6 samples were provided to the Dartmouth Genomics Shared Resource for sequencing. Viability of single-cell suspensions of was confirmed to be >90% using a Cellometer K2 (Nexcelom Bioscience). Samples were diluted to achieve a target cell capture of 10,000 cells and loaded on a Chromium Single Cell system (10X Genomics) to generate barcoded single-cell gel beads in emulsion, and scRNA-seq libraries were prepared using Single Cell 3′ Version 2 chemistry. Libraries were multiplexed and sequenced on 4 lanes of a Nextseq 500 sequencer (Illumina) with 3 sequencing runs. Demultiplexing and barcode processing of raw sequencing data was conducted using Cell Ranger v. 3.0.1 (10X Genomics). Reads were aligned to zebrafish (GRz11) to generate unique molecular index (UMI) count matrices.

### Data analyses

Count matrices produced from Cell Ranger were loaded into the R statistical working environment implemented on an R Studio programming platform (version 2023.12.1+402). Data quality analysis and filtering was conducted using scran (v1.30) as follows: First, the percentage of mitochondrial and hemoglobin transcripts (gene symbols prefixed by *mt-* and *hb-*respectively) were calculated per cell. Cells with < 500 or >60,000 unique molecular identifiers (UMIs: low quality and doublets), <200 or >6,000 features (low quality), or greater than 10% of reads mapped to mitochondrial genes (dying) were filtered from further analysis. Total cells per sample after filtering ranged from 809 to 2586. Data normalization, sample integration, dimension reduction and visualization was then conducted with Seurat (v5) using SCTransform^4^ and IntegrateData functions as implemented in Seurat v5^5^. Principal components were identified from the integrated dataset and were used for Uniform Manifold Approximation and Projection (UMAP) visualization of the data in 2D space. A shared nearest neighbor graph was constructed using default parameters, and clusters generated using the SLM algorithm in Seurat at a resolution of 0.5. The first 25 dimensions were used to identify 27 cell clusters (Table S1). Identities of individual cells were predicted using the scZCL package, which compares gene expression profiles of cells with Zebrafish Cell Landscape single cell data^6^,^7^ and dominant predicted identities for each cluster were identified. This resulted in a total of 14 predicted kidney marrow cell types in the dataset, ranging from 66 to 4294 cells total per cell type (Table S2, S3). Cells that were not labeled confidently were assigned “NA.” Statistical comparison of cell type proportion differences between genotypes were estimated using a 2-way ANOVA (cell type and genotype as factors), followed by Bonferroni’s multiple comparisons test within GraphPad Prism 9.0. Gene Ontology assessments were conducted for all differentially expressed genes within DAVID bioinformatics (https://david.ncifcrf.gov/). For differential gene expression within each cell type, single cell data was pseudobulked by sample and genewise exact tests for differences in the means between WT and AHR2null groups was conducted using EdgeR (padj <0.05).

## Results and discussion

### Absence of AHR2 Does Not Alter the Detailed Immune Cell Landscape

Based on cluster analysis, we were able to identify 14 cell types within our dataset across WT and AHR2null samples (Figure 1A). The mean percentage of cells per cell type ranged from 0.5 to 38.3 across the collective six samples, with erythrocytes constituting the highest (Figure 1B; Tables S3, S4). Within this paper, we focused on progenitor and mature immune cell types, excluding erythrocytes, thrombocytes or other kidney cells for our discussion and interpretation. Statistical analyses did not reveal any differences in proportions of progenitor or mature immune cell types between the two genotypes, indicating a lack of effect of AHR2 mutation on overall cell populations.

**Figure 1.**
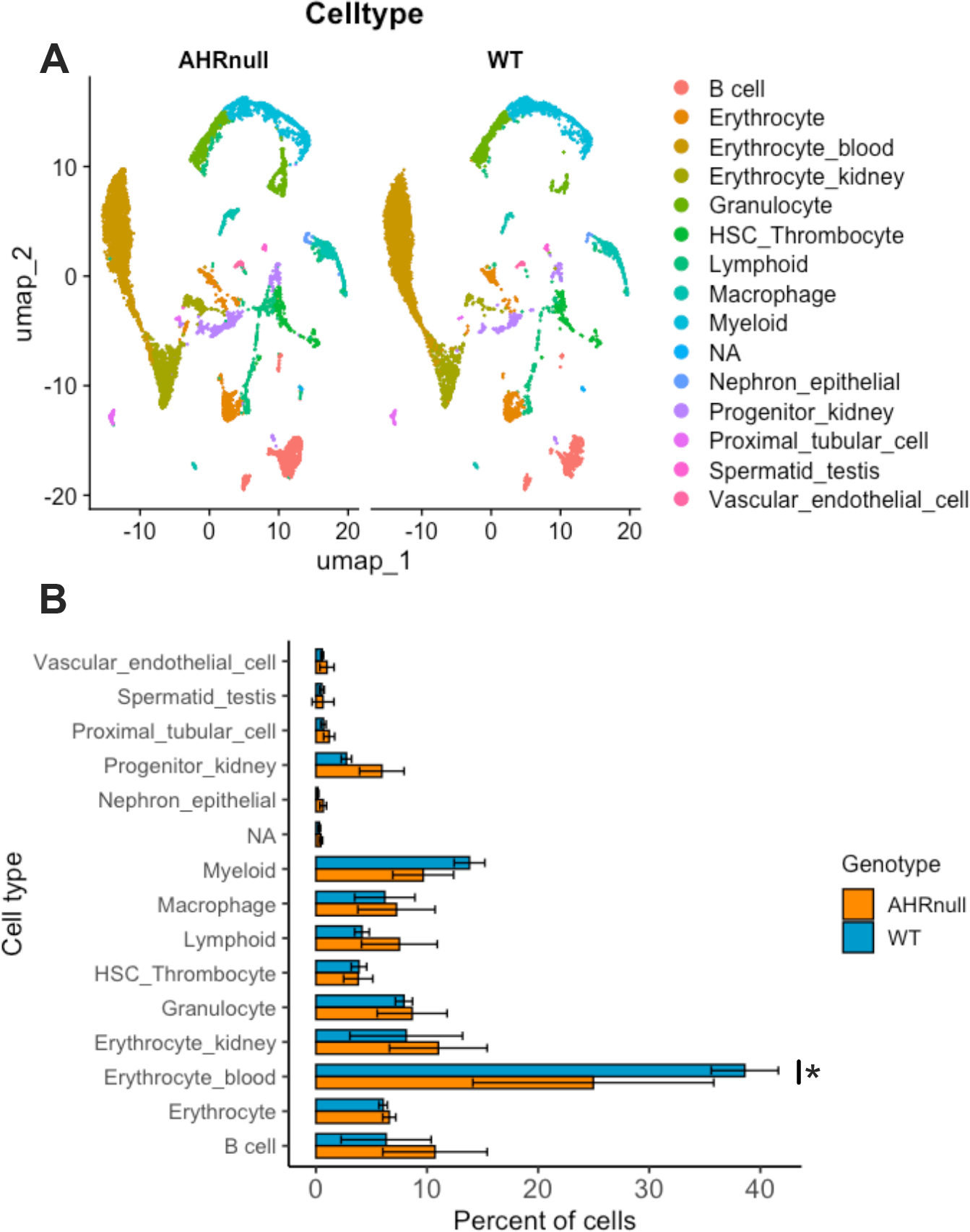
Cell types and proportion within wildtype and mutant kidney marrows. (A) UMAP plots of distribution of different cell types. (B) Comparison of cell type percentages between the two genotypes, expressed as mean of 3 replicates ± SD. * statistically significant difference using two-way ANOVA, followed by Bonferroni’s multiple comparisons test (p<0.05).

### Absence of AHR2 induces cell type-specific gene expression changes

Figure 2A presents a heat map of all genes that were dysregulated across various cell types, Figures 2 B-G presents UMAP plots of several genes discussed here and Table S5 lists differentially expressed genes. We first conducted gene ontology assessment of all differentially expressed genes within progenitor and mature immune cells (irrespective of cell types) between wildtype and AHR2null; this did not produce any significant results since the number of genes was low. At a gene level, we first focused on *ahr2* and *cyp1a*-constituents of the AHR pathway. Consistent with human *CYP1A1* datasets within the Human Protein Atlas (https://www.proteinatlas.org/), immune cells lacked expression of *cyp1a* (Figure 2B). However, expression of *ahr2* was present across multiple cell types but were not different within immune cells (Figures 2C, Table S5). We then focused on genes that were commonly dysregulated within multiple cell types. Examination of individual genes showed significantly high fold changes (>100 fold) for several genes, indicating that they have critical functions downstream of AHR2. Unfortunately, since many of the genes were unannotated or weakly annotated, it was not feasible to accurately assess function or gene ontology within the cell types. mRNA levels of *rrm2-*a cell cycle checkpoint gene^8^, were increased between 86 and 2980 times multiple immune cell types following AHR2 mutation (Figure 2D). Overexpression of *rrm2* has previously been implicated and identified as a biomarkers for increase in immune cell infiltration of tumor cells in several cancer types, including hepatocellular carcinoma ^9^. mRNA levels of si:ch211-181d7.3, predicted to regulate signal transduction, were increased within multiple immune cell types, including progenitor populations (∼120-800-fold changes; Figure 2E). Conversely, mRNA levels of *ndufaf5* (∼5-11-fold; Figure 2F), si:ch211-196c10.15 (∼4-5 fold) and si:rp71-36a1.1 (∼7-120-fold) were reduced within progenitors and mature immune cells. Among these, *ndufaf5* is known to be a regulator of mitochondrial chain complex^10^, indicating that AHR2 may directly or indirectly regulate mitochondrial function. Our data also revealed gene sets for which mRNA levels were uniquely altered within specific cell types. For example, markers of different carcinomas, such as *chl1a* (a cell adhesion protein molecule) and *tspan15* (a cell surface protein; Figure 2G), showed 4-8-fold changes in myeloid and macrophages respectively. Taken together, our data indicates that AHR2 potentially regulates several carcinogenic markers, primarily within progenitor populations and select mature immune cell types.

**Figure 2.**
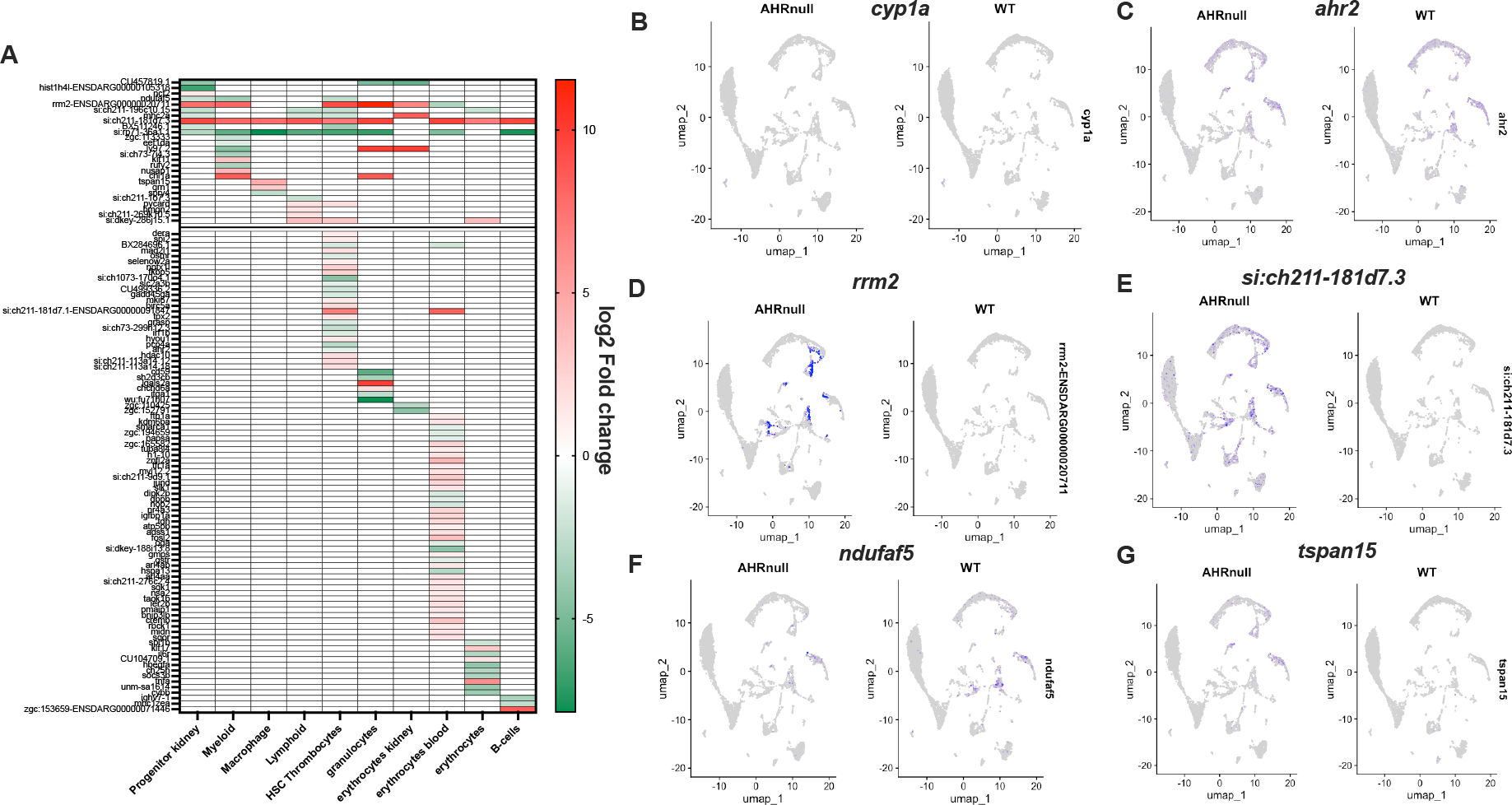
Gene expression differences within each cell type. (A) Heatmap of differentially expressed genes across cell types. (B-G) UMAP plots of representative genes between wildtype and mutant kidney marrows.

## Conclusion

Overall, our study shows several gene expression changes within immune and blood cells as a function of AHR2 mutation and many of these genes are identified as biomarkers of cancer. While many single cell studies use 1 or 2 biological replicates owing to the cost of the technique, we conducted our study with 3 replicates per genotype for robust statistics. Unfortunately, lack of annotation, coupled with lack of functional information limited us from through investigation of AHR2 regulation of hematopoiesis. Several differentially expressed, but unannotated genes showed exceedingly high fold changes in AHR2 mutants, highlighting their importance in AHR2 regulation of hematopoiesis. Therefore, proper characterizations of these genes, coupled with functional studies are required. Despite these shortcomings, this work shows that AHR2 is important in endogenous genetic regulation within several immune cell types and creates a platform for in-depth exploration of these targets.

## Acknowledgements

The authors thank Fred Kolling, the Dartmouth Cancer Center Shared Genomics Resource and the Leach Laboratory for single cell sequencing support.

